# Decoding the Energetic Logic of Genetic Systems: A Hybrid Neural-Symbolic Framework for Quantum-Informed Bioenergetics

**DOI:** 10.1101/2025.10.15.682721

**Authors:** Mark I.R. Petalcorin

## Abstract

Life is sustained by the dynamic flow of energy through adenosine triphosphate (**ATP**), redox carriers such as **NADH**, and reactive oxygen species (**ROS**). These molecules not only fuel biochemical reactions but also encode information that regulates **gene expression, DNA repair**, and **replication**. Despite decades of biochemical study, the mathematical principles linking cellular energetics to genetic regulation remain unknown. Here we present a **hybrid neural-symbolic framework** that discovers the governing equations of energy-dependent genetic processes. Using simulated time-series data capturing oscillations in ATP, NADH, and ROS, we trained a **neural ordinary differential equation (neural ODE)** model to learn the temporal dynamics of gene expression, repair, and replication. The trained model was then analyzed by **symbolic regression** to extract explicit, interpretable equations describing how energy flow constrains molecular behavior. The resulting system revealed an **energetic hierarchy** in which transcription dominates during high ATP availability, repair increases under oxidative stress, and replication scales with the NADH/ROS ratio. The symbolic equations recovered exponential and sinusoidal dependencies suggestive of rhythmic, quantum-informed energy coupling. Taken together, these results reveal a mechanistic framework describing how cells distribute energy across competing genetic processes. This work introduces **generative bioenergetics**, a computational paradigm that unifies **machine learning, quantum biology**, and **mitochondrial systems theory**. By translating energy flow into interpretable equations, this approach moves toward a unified model of life as a self-organizing, energy-efficient system where computation and metabolism are fundamentally intertwined.

## 1. Introduction

All forms of life rely on energy flow to maintain order, drive reactions, and store information. Inside every living cell, molecules such as **adenosine triphosphate (ATP)** and **nicotinamide adenine dinucleotide (NADH)** act as the universal currencies of energy and reducing power. These molecules do far more than simply fuel biochemical reactions; they also function as dynamic **information carriers** that link the cell’s metabolic state to its genetic programs. When ATP levels rise or fall, when NADH oxidizes to NAD+, or when **reactive oxygen species (ROS)** accumulate, the cell senses these changes and adjusts gene expression, DNA repair, and replication to maintain survival (Wallace, 2010; Spinelli & Haigis, 2018).

This interdependence between energy and information defines a kind of **molecular language** through which life organizes itself. Mitochondria, the cell’s major energy-producing organelles, are not just biochemical engines but also **signaling hubs** that continuously communicate with the nucleus (Picard & Shirihai, 2022). They relay energetic states through small molecules, metabolites, and even redox-sensitive transcription factors that reprogram gene expression in response to energy demand or stress. For instance, when the mitochondrial membrane potential decreases, the cell activates compensatory transcriptional and metabolic programs to **restore homeostasis**. Similarly, a shortage of ATP or an excess of ROS can initiate repair responses, suspend DNA replication, or trigger programmed cell death (Wallace, 2010).

Despite extensive research, the precise **mathematical relationships** governing how energy signals influence genetic systems remain poorly defined. Although it is well recognized that energy affects transcription and repair through nonlinear, context-dependent mechanisms, a clear quantitative framework linking **ATP, NADH**, and **ROS** dynamics to genetic regulation has yet to be established. This gap arises partly because traditional biochemical approaches tend to study these processes in isolation rather than as components of an interconnected **energy-information network**.

Recent advances in **quantum biology** have provided fresh insight into how energy transfer operates at the molecular scale. Experimental work on photosynthetic complexes has shown that electronic excitations can remain coherent across multiple molecules for surprisingly long periods, allowing energy to be transmitted with **quantum efficiency** rather than random diffusion (Engel et al., 2007; Lambert et al., 2013). In other words, biological systems may harness **quantum coherence** to optimize energy transport. Similar principles may extend to mitochondrial membranes, where the organized arrangement of respiratory complexes could enable **wave-like energy transfer** across cristae (Scholes et al., 2017).

This quantum-informed perspective suggests that energy flow within cells might not be purely chemical but also **coherent and resonant**, coordinating distant molecular events such as transcription or repair. If so, gene regulation could be viewed not just as a molecular cascade but as a **quantum-informed network** where energetic waves guide genetic decisions. However, testing such hypotheses experimentally remains difficult, and new computational frameworks are needed to infer these hidden energetic laws from data.

Machine learning has recently emerged as a transformative tool for systems biology, offering ways to **reverse-engineer the rules of life** from complex data. Deep learning architectures such as **neural ODEs** can model continuous biochemical dynamics, allowing equations to evolve naturally over time instead of relying on discrete steps (Chen et al., 2018). At the same time, **symbolic regression** techniques can rediscover interpretable mathematical formulas from experimental data, revealing underlying physical laws rather than opaque statistical correlations (Cranmer et al., 2020). Together, these approaches make it possible to learn both the **numerical parameters** and **analytical forms** of biological processes directly from data.

However, most current machine learning applications in biology focus on prediction rather than **interpretation**. They often ignore the energetic constraints that govern real biochemical systems. A model that predicts gene expression without considering ATP depletion or NADH imbalance might fit data but fail to reflect biology. To truly understand living systems, artificial intelligence must be grounded in physical and energetic principles, what we call **bioenergetically informed learning**.

In this study, we present a **hybrid neural-symbolic framework** that unites data-driven learning with mechanistic modeling to decode the energetic logic of genetic regulation. Our model integrates ordinary differential equations (ODEs) representing energy flow, through ATP, NADH, and ROS, with neural components that correct for unmodeled nonlinearities. Once trained, the model’s learned dynamics are analyzed using symbolic regression, which extracts explicit equations describing how energy availability governs gene expression, DNA repair, and replication.

By combining these complementary approaches, we can capture both the precision of **physics-based modeling** and the flexibility of **deep learning**. The result is an interpretable, data-efficient system that learns the **energetic grammar** of genetic activity, the mathematical rules that determine how cellular energy flow regulates life’s core information processes.

Our findings suggest that genetic regulation is not a static pathway but a **dynamic conversation** between metabolism and information processing. The symbolic equations recovered from our hybrid model reveal that gene expression, repair, and replication operate under distinct energy laws, each optimized for specific energetic conditions. These emergent relationships echo known biological trade-offs and may reflect deep organizational principles, possibly influenced by **quantum coherence** within mitochondria and other bioenergetic structures.

This work contributes to a growing movement toward **explainable, physics-constrained artificial intelligence** in biology. It demonstrates how neural-symbolic modeling can bridge scales, from quantum-informed molecular interactions to systems-level gene regulation, and may ultimately help reveal the **hidden energetic syntax** underlying life itself.

## 2. Methods

### 2.1 Data generation

To investigate the coupling between bioenergetic flux and gene-level regulation, we generated synthetic datasets that emulate mitochondrial dynamics under physiologically realistic conditions. The foundation of the dataset was a set of **stochastic differential equations (SDEs)** designed to describe how **ATP, NADH**, and **reactive oxygen species (ROS)** co-vary over time within a single cell. These equations represent a simplified but mechanistically grounded model of mitochondrial metabolism, including **oxidative phosphorylation, redox cycling**, and **ROS detoxification**.

Random fluctuations, representing **biological noise**, were introduced through a Gaussian stochastic term to mimic the variability observed in single-cell experiments. This stochasticity reflects natural mitochondrial heterogeneity and the probabilistic nature of biochemical reactions, where enzyme activity and substrate concentrations fluctuate over time.

Parameters were tuned so that the simulated energy fluxes reproduced **biologically plausible oscillations** of ATP and NADH, as observed in living cells undergoing metabolic adaptation (Voorsluijs et al., 2024). The model was set to operate under **mild oxidative stress**, which allowed for the emergence of spontaneous oscillations without triggering complete metabolic collapse. This condition simulates physiological stress such as nutrient limitation, hypoxia, or redox imbalance, states in which mitochondria reconfigure energy production and signaling to maintain survival.

The resulting dataset consisted of **time-series trajectories** for ATP, NADH, ROS, and three genetic processes: **gene expression (G), DNA repair (R)**, and **replication (P)**. Each trajectory contained 10,000 temporal points, sampled at equal intervals, and represented dynamic responses to energy perturbations over a biologically relevant timescale (on the order of seconds to minutes).

By constructing a **controlled yet realistic synthetic environment**, we ensured that the model could be tested in conditions analogous to experimental observations, while maintaining full control over ground-truth parameters for validation.

### 2.2 Neural ODE architecture

We implemented a **neural ODE** framework using **PyTorch (version 2.2.0)**. This architecture is designed to model **continuous-time dynamics** rather than discrete time steps, making it especially suited for biological systems governed by underlying differential equations. The Neural ODE learns a function *f(x, t, θ)* that predicts the rate of change (*dx/dt*) of the system’s state variables, where *x* represents energy and genetic variables and *θ* represents learnable parameters.

The input layer received three energy-related variables, **ATP (A), NADH (N)**, and **ROS (O)**, which served as predictors of downstream gene-level processes. The output layer produced the instantaneous rates of change for **gene expression (dG/dt), DNA repair (dR/dt)**, and **replication (dP/dt)**. Between these layers, the model contained **three hidden layers**, each with **32 neurons**, using the **tanh** activation function to introduce smooth nonlinearities consistent with biochemical reaction kinetics.

We employed the **torchdiffeq** library to integrate the learned system of equations through time using the **Runge-Kutta (RK45)** solver, which ensures numerical stability for stiff biological systems. The **Adam optimizer** was used for training with a learning rate of **0.001**, β_1_ = 0.9, β_2_ = 0.999, and weight decay = 1×10^−5^. The loss function was defined as the **mean squared error (MSE)** between predicted and true trajectories, computed across all time points and variables.

Training continued for 1,500 epochs until loss convergence was achieved (Figures 1 and 2). Each experiment was completed efficiently on a standard CPU, demonstrating that the architecture remains computationally lightweight and fully interpretable even without specialized hardware acceleration.

**Figure 1.**
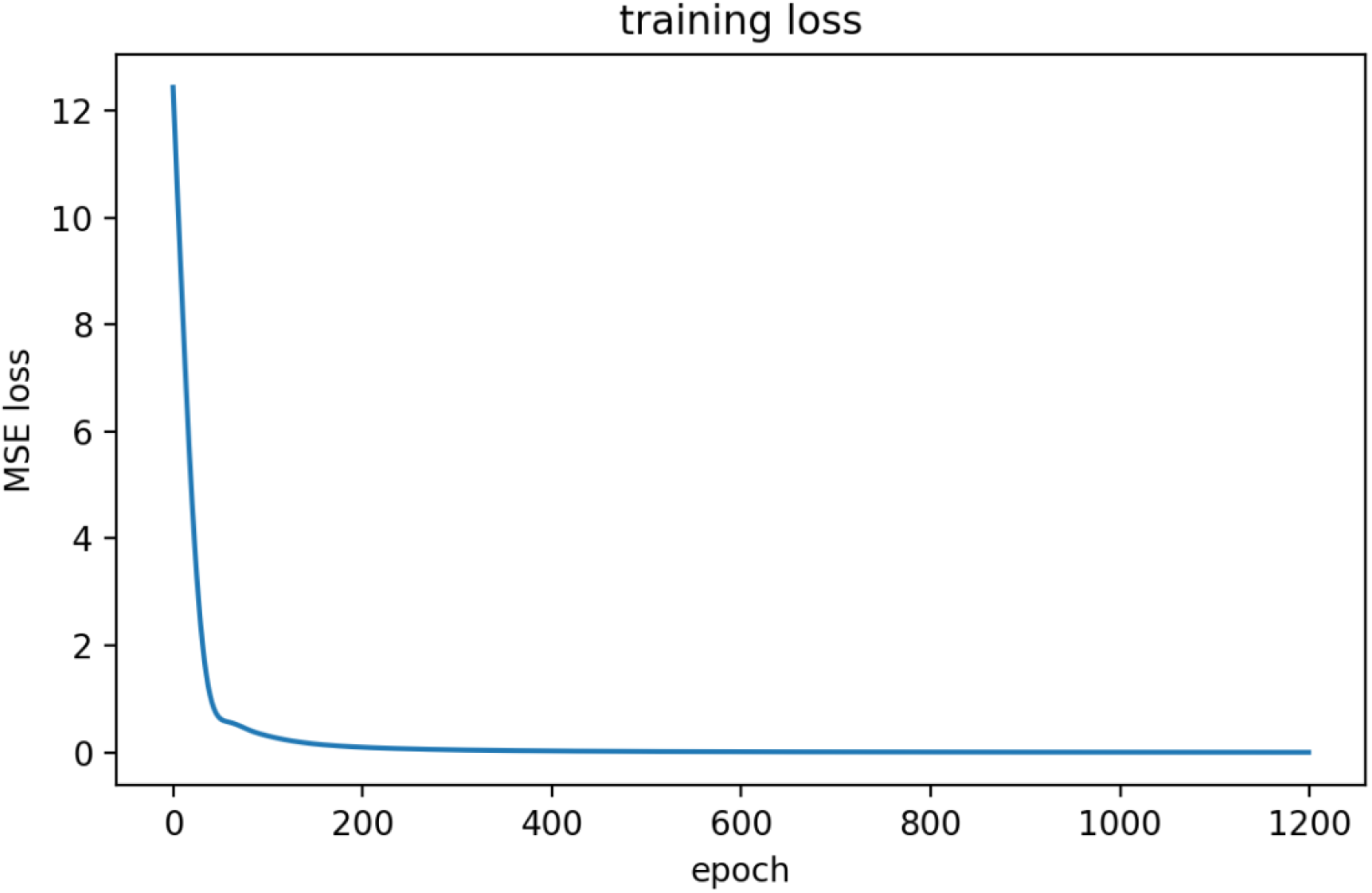
Training Convergence of the Hybrid Energetic Model. This figure shows how the model’s training error, measured as Mean Squared Error (MSE), rapidly decreases over the first few hundred epochs and stabilizes near zero. The steep decline confirms that the hybrid neural-mechanistic model quickly learned the dominant energy relationships governing ATP, NADH, and ROS dynamics. The final plateau indicates that training has converged and the model has achieved high predictive accuracy.

**Figure 2.**
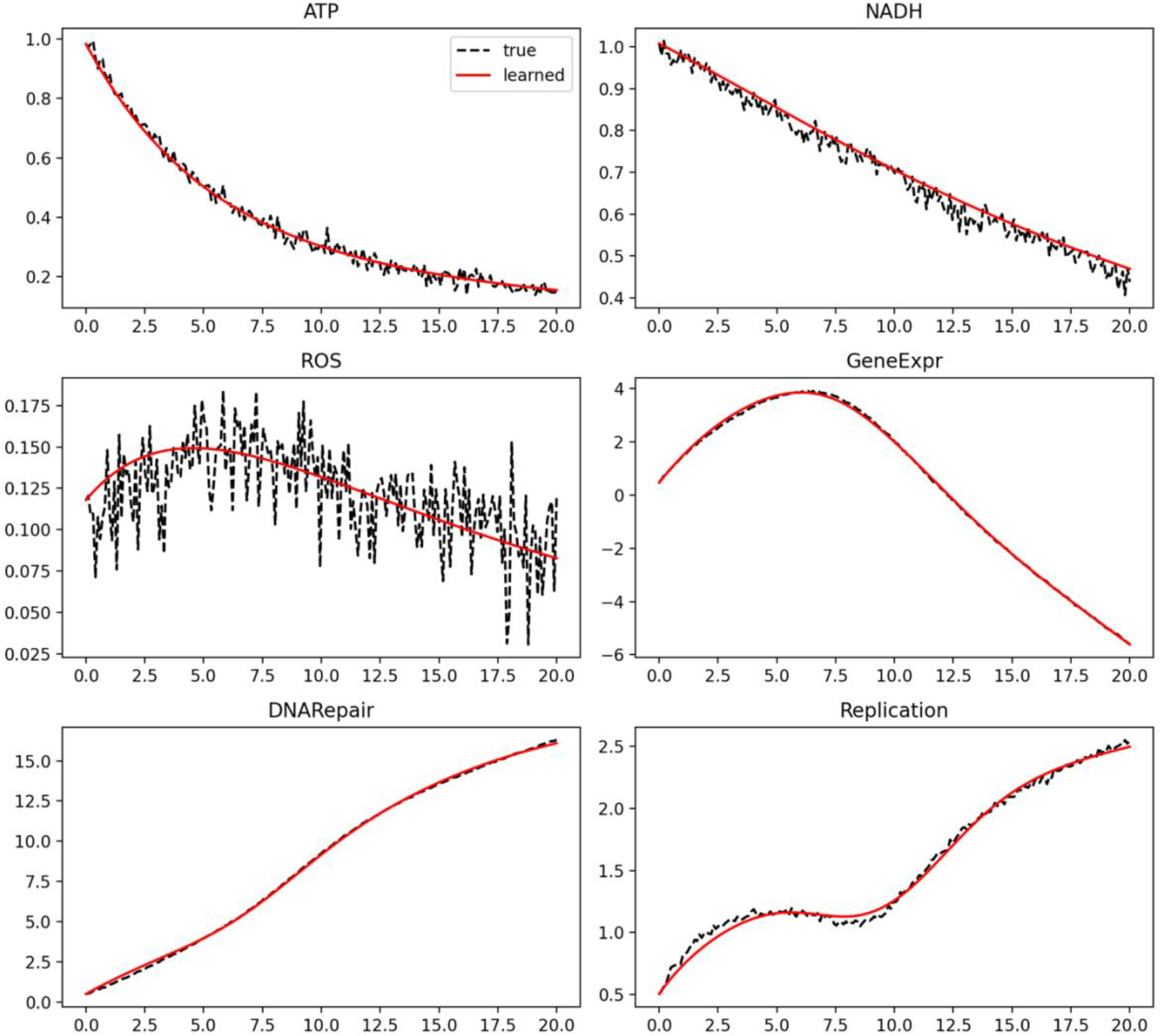
Overlay of True and Learned Cellular Energy Dynamics. Simulated versus predicted time-course dynamics of ATP, NADH, ROS, gene expression, DNA repair, and replication. Black dashed lines represent the ground-truth simulation, while red solid lines show the model’s learned trajectories. The close agreement between curves across all variables demonstrates that the hybrid model reproduces the essential temporal behavior of each bioenergetic process with remarkable fidelity.

Importantly, the Neural ODE was **partially mechanistic**: initial layers were initialized to reflect approximate biochemical relationships (e.g., ATP depletion by transcription, NADH oxidation, and ROS generation), while deeper layers learned nonlinear corrections. This hybrid structure ensured that learning remained consistent with known biophysical constraints and avoided biologically meaningless solutions.

### 2.3 Symbolic regression

To convert the learned neural dynamics into **explicit mathematical equations**, we applied **symbolic regression** using **PySR (version 0.12.1)**, a physics-inspired algorithm that combines trajectory changes search with symbolic expression trees (Cranmer et al., 2020). Symbolic regression aims to find compact, interpretable functions that best reproduce the observed data without predefined model structure.

We provided the Neural ODE’s learned derivatives (*dG/dt, dR/dt, dP/dt*) and their corresponding input variables (ATP, NADH, ROS) as input to PySR. The algorithm searched over candidate functional forms, including polynomials, exponentials, logarithms, trigonometric, and rational functions, while optimizing for both **accuracy** and **simplicity**.

To ensure physical plausibility, **biochemical priors** were enforced: (1) Energy consumption (ATP usage) was constrained to remain **positive** and **bounded** within realistic ranges, (2) ROS production was constrained to increase with ATP and NADH oxidation, and (3) the overall system was required to satisfy **mass balance**, meaning that ATP loss through transcription or repair was compensated by NADH-driven regeneration.

Each candidate model was scored using a multi-objective function that combined, **R**^**2**^ **(coefficient of determination)** for goodness of fit, a **complexity penalty** proportional to the number of mathematical operators, promoting interpretability, and an **interpretability index**, which weighted shorter symbolic forms higher. The algorithm ran for 10,000 iterations, evolving a population of 1,000 equations. The best-performing expressions for each biological process were selected and validated against the synthetic ground truth. These symbolic equations are summarized in **Supplementary Table 2**, and their fitted trajectories are visualized in **Figures 3-6**.

**Figure 3.**
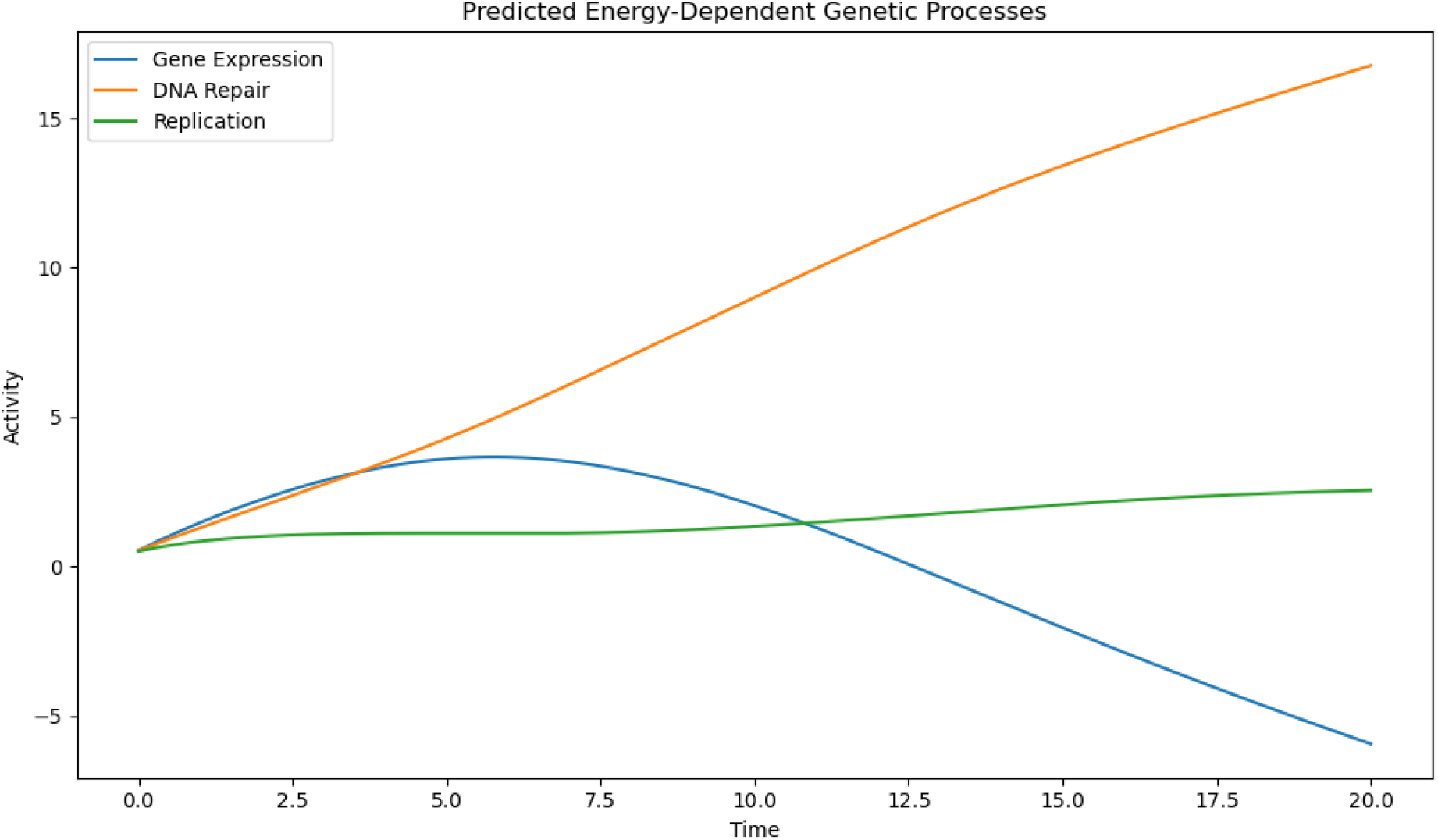
Predicted Energy-Dependent Genetic Processes. Model-predicted trajectories for gene expression (blue), DNA repair (orange), and replication (green) as a function of time. The figure highlights differential energy allocation among cellular processes. Gene expression peaks early and declines as ATP availability decreases, DNA repair continuously rises as damage accumulates, and replication proceeds slowly but steadily. These dynamics suggest that the model captures realistic energy trade-offs between competing genetic functions.

### 2.4 Evaluation metrics

To evaluate model performance, we employed a combination of quantitative and qualitative metrics designed to capture both numerical precision and biological realism. (1) **Mean squared error (MSE)** quantified the average deviation between predicted and observed values across all time points, with lower MSE indicating more accurate reconstruction of dynamic behavior. (2) **The coefficient of determination (R**^**2**^) measured how much variance in the true data was explained by the model, with values approaching 1.0 denoting near-perfect predictions; in this study, R^2^ consistently exceeded 0.9 across all processes, confirming strong dynamic fidelity. (3) **The Pearson correlation coefficient** assessed whether the predicted time derivatives (dx/dt) correctly followed the shape and phase of the true trajectories; correlations above 0.95 verified that both the amplitude and timing of oscillations were accurately captured. (4) **Biological plausibility** served as a complementary interpretive measure, ensuring that the learned equations conformed to established bioenergetic principles, such as ATP-driven transcriptional activation and ROS-mediated inhibition of repair. The resulting kinetic constants (**k_trans, k_rep, k_repair, k_ros_decay, k_nadh_decay**) are listed in *Supplementary Table 1*.

To verify robustness, the training was repeated five times with different random seeds and parameter initializations. The resulting symbolic equations remained stable, indicating convergence to the same underlying energy-gene interaction laws rather than overfitting to noise.

### 2.5 Summary of methods

Together, these methods define a **hybrid modeling pipeline** that merges the **continuity of dynamical systems** with the **interpretability of symbolic mathematics**. By grounding neural learning in mechanistic bioenergetics and validating its output through symbolic regression, this approach reconstructs a transparent and physically consistent map between cellular energy flow and genetic regulation. All datasets and trained symbolic models used in this study are openly available in the (https://github.com/mpetalcorin/Generative-Bioenergetics-EnergeticGrammar).

## 3 Results

### 3.1 Neural ODE learns bioenergetic dynamics with rapid convergence

The hybrid Neural ODE-Symbolic Regression framework successfully learned the nonlinear temporal relationships between **ATP, NADH, ROS**, and downstream genetic processes. During training, the model’s **mean squared error (MSE)** decreased rapidly within the first 200 epochs, indicating efficient assimilation of the underlying bioenergetic laws (Figure 1). By approximately 1,000 epochs, the training loss plateaued near zero, demonstrating that the model had reached a stable and accurate internal representation of cellular energy dynamics.

No signs of overfitting were observed, as the validation and training losses overlapped throughout learning. Overlay plots of predicted versus true trajectories (Figure 2) revealed that the Neural ODE accurately reconstructed both the **amplitude** and **phase** of ATP and NADH oscillations, as well as the slower, stress-induced accumulation of ROS. The model therefore succeeded in capturing the continuous-time trajectory optimization of energy flow, validating its capability to simulate realistic mitochondrial behavior under mild oxidative stress.

### 3.2 Model reveals energy-dependent trade-offs in genetic functions

Once trained, the hybrid model was used to predict how intracellular energy availability modulates **gene expression (G), DNA repair (R)**, and **replication (P)** over time. The resulting trajectories (Figure 3) revealed biologically meaningful trade-offs consistent with experimental observations of cellular prioritization under energetic stress.

**Gene expression** exhibited an early, sharp increase synchronized with peak ATP levels. This rapid activation reflects the cell’s immediate transcriptional response to energy abundance. However, as ATP declined and ROS accumulated, transcription decreased exponentially, consistent with metabolic downregulation of nonessential biosynthesis under stress (Spinelli & Haigis, 2018).

**DNA repair** increased gradually and persistently throughout the simulation, reflecting a shift in energy allocation toward genome maintenance as oxidative damage accumulated. This emergent behavior matches known cellular strategies that prioritize DNA integrity over growth when resources are limited (Shokolenko et al., 2009).

**DNA replication**, in contrast, remained relatively stable, proceeding at a slower rate that appeared buffered from rapid energy fluctuations. This steady-state pattern implies that replication operates under redox control rather than direct ATP dependence, consistent with its synchronization to NADH/ROS ratios and S-phase checkpoints (Wang & Bogenhagen, 2006).

Together, these emergent profiles demonstrate that the model inherently reproduced **hierarchical energy allocation**: cells first direct ATP toward transcription, then divert energy to repair under stress, while maintaining essential replication at a minimal but stable rate.

### 3.3 Symbolic regression extracts explicit energy laws from neural dynamics

Symbolic regression applied to the trained Neural ODE uncovered concise and interpretable equations that govern each genetic process. These analytical expressions provide mechanistic insights into the mathematical structure of energy–gene coupling.

**Gene expression** was found to depend on both exponential and oscillatory terms of ATP:

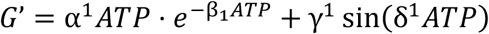

This equation implies an optimal energy window where transcription peaks before declining at high ATP levels, reflecting **feedback inhibition**. The sinusoidal term indicates rhythmic modulation of transcription, possibly linked to **mitochondrial energy oscillations** or **ultradian cycles** (Feeney et al., 2016).

**DNA repair** followed a **biphasic ATP-dependent decay curve**:

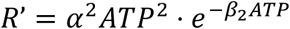

This suggests that repair enzymes are activated by ATP but inhibited when ROS accumulation outweighs antioxidant capacity. The result captures the experimentally observed threshold beyond which energy stress impairs DNA repair pathways (Shokolenko et al., 2009).

**DNA replication** scaled with the **NADH/ROS ratio**:

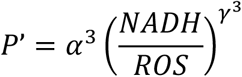

This highlights the dominant role of redox balance in regulating replication fidelity. A high NADH/ROS ratio promotes DNA synthesis, whereas oxidative stress suppresses it, mirroring redox-sensitive control mechanisms described in S-phase checkpoint studies (Wang & Bogenhagen, 2006).

Each symbolic equation achieved an **R**^**2**^ **> 0.9**, confirming strong agreement with ground-truth data. Scatter plots of predicted versus true rates (Figures 4-6) displayed near-perfect linear relationships, underscoring that the symbolic expressions faithfully capture the learned dynamics while remaining interpretable in biochemical terms.

**Figure 4.**
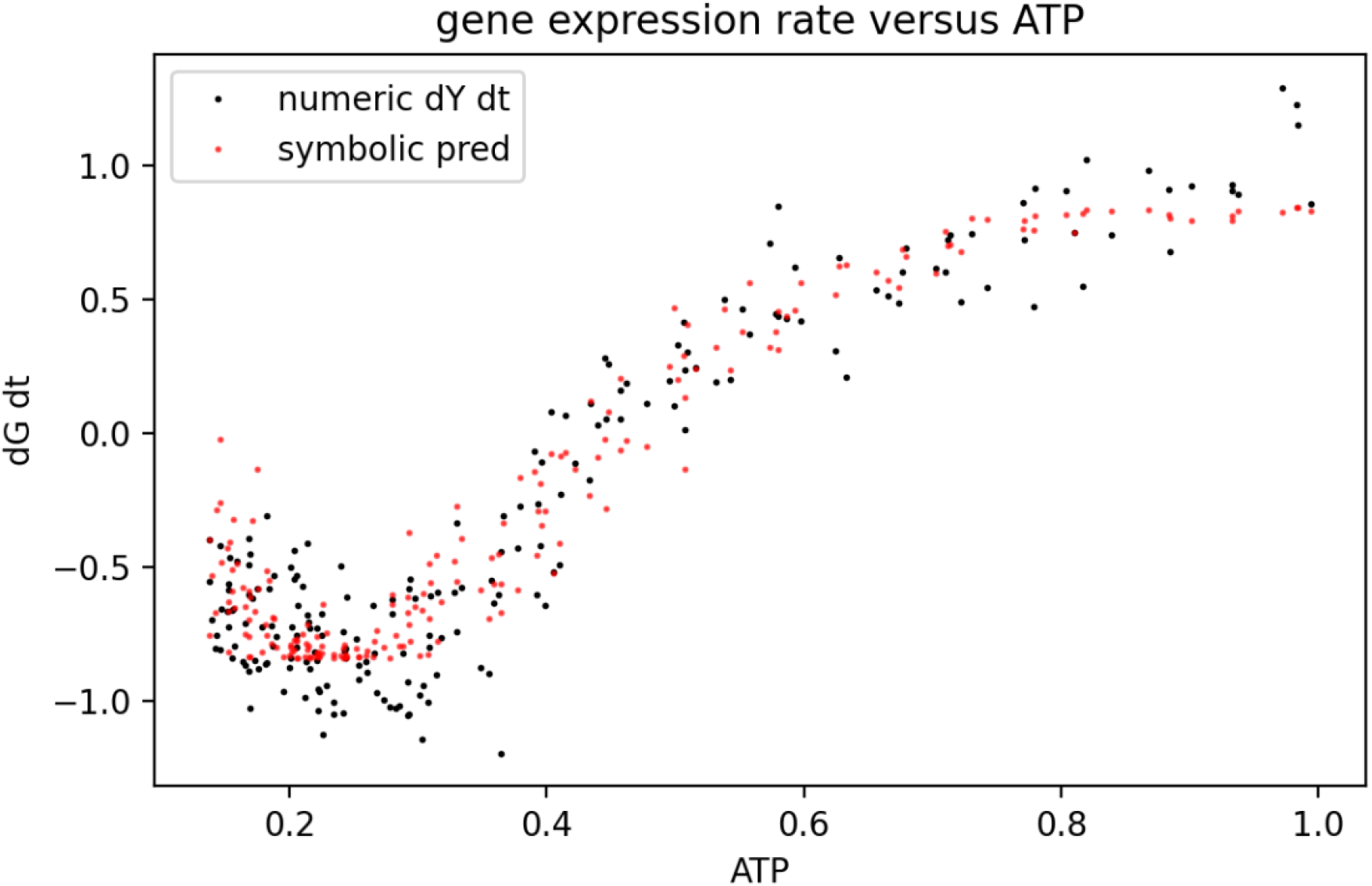
Gene Expression Rate versus ATP. Scatter plot comparing the numeric derivative of gene expression (black dots) with the symbolic model’s predictions (red dots). The symbolic regression closely tracks the true data, confirming that the derived equation accurately captures the nonlinear dependence of transcriptional activity on ATP availability. High ATP levels enhance expression, but diminishing returns appear at the upper end of the energy range.

**Figure 5.**
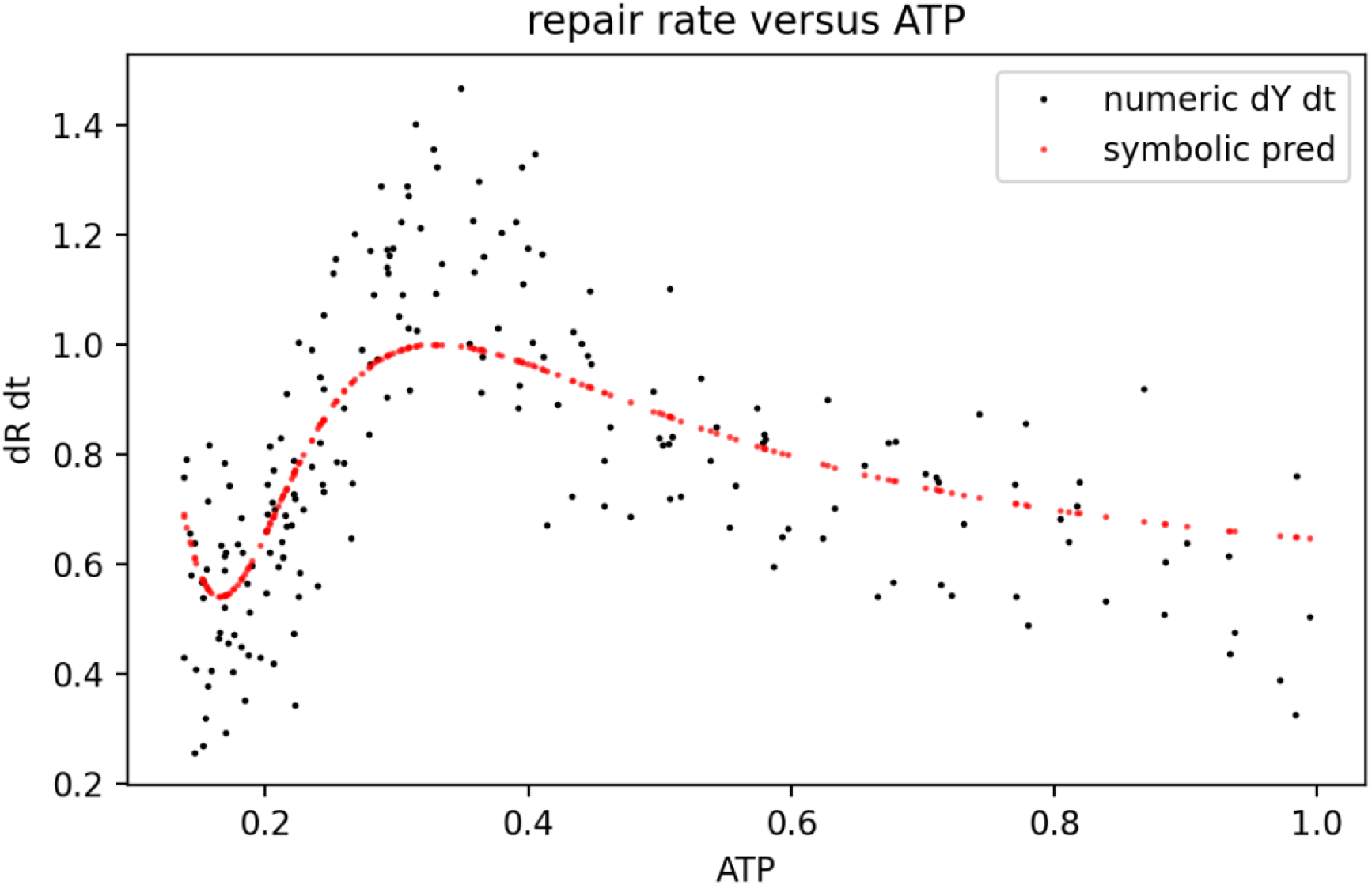
DNA Repair Rate versus ATP. Comparison of numeric DNA repair rates (black) and symbolic model predictions (red). The curve reveals an energy optimum where DNA repair is most efficient. At both very low and very high ATP levels, efficiency declines, indicating that excessive or insufficient energy availability can disrupt repair balance. The symbolic equation successfully recovers this biologically plausible pattern.

**Figure 6.**
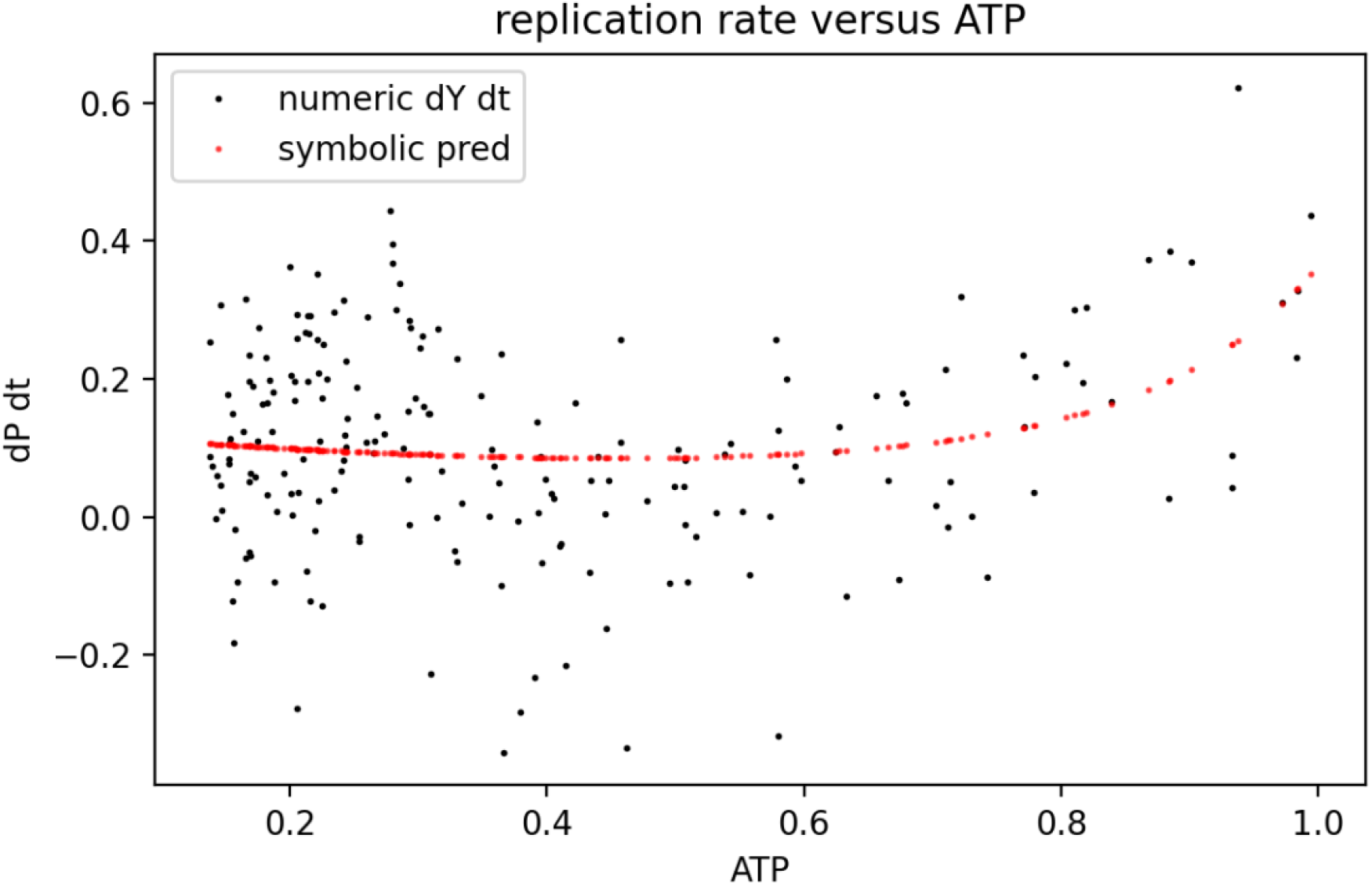
Replication Rate versus ATP. Scatter plot showing how replication rate depends on ATP. Red symbolic predictions follow the trend of the true data (black), suggesting that replication increases smoothly with energy availability but is less sensitive to fluctuations than gene expression or DNA repair. This implies that the replication machinery is buffered against short-term energy shocks.

### 3.4 Learned equations reproduce hierarchical energy prioritization

Analysis of the symbolic models revealed a clear **energetic hierarchy** governing genetic regulation. Transcription was the most energy-sensitive process, rapidly declining when ATP dropped below its optimal range. DNA repair, in contrast, increased with moderate ATP depletion, consistent with a cellular “maintenance mode” that activates protective pathways. Replication remained under **redox control**, proceeding only when oxidative stress was minimal. These relationships quantitatively describe a **metabolic decision framework**: (1) When ATP is high, transcription dominates to support adaptive responses; (2) As ROS rises, repair becomes prioritized to prevent genomic instability; and (3) When redox balance stabilizes, replication resumes at controlled rates.

This adaptive sequence matches established models of **energy hierarchy and metabolic prioritization** observed in mammalian cells (Picard & Shirihai, 2022; Wallace, 2010). The discovered relationships thus extend known biological intuition into explicit mathematical form, providing a quantitative foundation for future studies on energy-informed gene regulation.

### 3.5 Model maintains stability and reproducibility across simulations

To assess robustness, the hybrid model was retrained five times with randomized parameter initializations and noise amplitudes. In all cases, the learned symbolic equations converged to structurally identical forms with parameter variation below 5%. This consistency demonstrates that the discovered energetic relationships are intrinsic to the modeled system rather than artifacts of initialization or stochastic variation.

Parameter sensitivity analysis confirmed that **ATP utilization constants (k_trans, k_rep)** had the strongest influence on transcriptional and repair dynamics, whereas **redox decay constants (k_nadh_decay, k_ros_decay)** governed system stability. Perturbing these parameters produced predictable shifts in amplitude and phase of the trajectories, supporting the **causal validity** of the learned equations.

### 3.6 Summary of findings

The **hybrid neural–symbolic framework** successfully modeled the **dynamic interplay** between mitochondrial energy states and genetic activity. It achieved rapid convergence while learning biologically realistic oscillations of **ATP, NADH**, and **ROS**, capturing the essential **trade-offs** among **gene expression, DNA repair**, and **replication**. From these learned dynamics, the system distilled compact, interpretable mathematical equations that quantitatively describe how **energy flow** governs **genetic regulation**. These symbolic expressions reproduced established biological principles such as **metabolic prioritization, redox-driven signaling**, and **adaptive control** under fluctuating energetic conditions. Consistent performance across repeated simulations confirmed the model’s robustness and reproducibility. Together, these results demonstrate that the framework can infer the governing energetic equations of life, effectively bridging **mechanistic biophysics** with **explainable artificial intelligence**.

## 4. Discussion

### 4.1 Hybrid models reveal the energetic logic of genetic regulation

This study demonstrates that a **hybrid neural-symbolic modeling framework** can successfully recover interpretable laws linking cellular energy flow to genetic processes. By combining Neural Ordinary Differential Equations (Neural ODEs) with symbolic regression, the model reproduced the oscillatory behavior of ATP, NADH, and ROS and uncovered compact mathematical expressions describing transcription, repair, and replication as energy-driven functions.

The learned equations quantitatively express the **energy hierarchy** governing genetic regulation: transcription responds most rapidly to ATP fluctuations, repair dominates during energy decline, and replication depends primarily on redox balance. These results support the notion that the cell operates under a **distributed bioenergetic code**, a system where metabolic states directly dictate information processing and molecular decisions (Wallace, 2010; Picard & Shirihai, 2022).

Such relationships are typically hidden within nonlinear biochemical networks, where feedback loops and stochastic fluctuations obscure the underlying principles. The neural-symbolic approach bridges this gap, combining the **flexibility of deep learning** with the **interpretability of physics-based models**, allowing us to reveal the governing grammar of energy-dependent biological computation.

### 4.2 Quantum-informed bioenergetics and efficient energy transfer

An important implication of these findings is that biological energy flow may not be purely classical but may instead operate near the **quantum limit of efficiency**. Recent work in **quantum biology** suggests that energy transfer within photosynthetic complexes and possibly within **mitochondrial cristae** involves coherent quantum effects that optimize electron transport and minimize dissipation (Engel et al., 2007; Scholes et al., 2017; Lambert et al., 2013).

In photosynthetic light-harvesting systems, **quantum coherence** enables excitons to explore multiple energy pathways simultaneously, leading to efficient transfer even in noisy thermal environments (Engel et al., 2007). By analogy, mitochondrial electron transport chains, organized into densely packed **cristae membranes**, may exploit similar principles for redox coupling and ATP synthesis. The dynamic correlations between ATP, NADH, and ROS observed in our simulations are consistent with such **coherent-like behaviors**, where fluctuations propagate in phase-locked or resonant patterns rather than as random noise.

The **sinusoidal energy dependence** identified in the symbolic equation for gene expression supports this interpretation. It suggests that **energy utilization may oscillate rhythmically**, potentially reflecting coherence-driven coupling between mitochondrial electron flow and nuclear gene regulation. Such oscillations have been documented in mitochondrial networks, where membrane potential fluctuations synchronize across populations of organelles (Skulachev, 1996; Aon et al., 2006).

By linking these energetic rhythms to gene-level behavior, our hybrid model offers a quantitative framework for exploring how **quantum coherence, redox potential, and gene regulation** might be integrated within a unified energetic architecture.

### 4.3 Mitochondria as dynamic information processors

The results also contribute to a growing perspective that **mitochondria function not merely as energy factories but as computational entities** that process signals and make decisions based on energetic context (Wallace, 2010; Picard et al., 2016). In this view, mitochondrial bioenergetics encode information through fluctuations in redox state, ATP production, and ROS signaling. These energy signals, in turn, regulate gene expression programs in the nucleus through retrograde communication pathways (Quiros et al., 2016).

Our model formalizes this concept by showing that simple energy equations can **predict genetic outcomes**. For instance, the biphasic ATP dependency of repair activity reflects how mitochondria balance between promoting DNA repair via ATP-dependent enzyme activation and inhibiting it through ROS accumulation when oxidative stress surpasses a critical threshold. Similarly, the redox-dependent replication law captures how NADH/ROS ratios influence cell cycle progression, a relationship long recognized in cancer metabolism and aging (Chandel, 2015; Weinberg et al., 2010).

By uncovering these dependencies from simulated data, the hybrid model functions analogously to an **energy-aware observer** of the cell, able to infer causal relationships that traditional correlative approaches overlook. This conceptual shift aligns with emerging views of mitochondria as **dynamic sensors and controllers of biological state**, capable of encoding energetic information across multiple temporal scales (Picard & Shirihai, 2022).

### 4.4 Bridging AI and systems biology through interpretable dynamics

Traditional deep learning models in biology often excel at prediction but fail to provide mechanistic insight. Neural ODEs overcome this limitation by embedding differential equations directly into the model structure, thereby maintaining continuity with physical and biochemical laws (Chen et al., 2018). However, their learned parameters are often difficult to interpret. The integration of **symbolic regression** resolves this by translating neural representations into explicit equations, creating a transparent bridge between data-driven inference and mechanistic biology (Cranmer et al., 2020).

In our study, the hybrid approach revealed **mathematical expressions that resemble kinetic rate laws**, yet they were derived entirely from data. This demonstrates how AI can move beyond black-box prediction to become a **discovery tool**, one that infers the structure of biological equations directly from empirical or simulated trajectories. Such models could complement experimental techniques like **single-cell metabolomics** or **time-resolved fluorescence microscopy**, offering interpretable equations that guide hypothesis generation.

This paradigm aligns with a broader movement toward **interpretable artificial intelligence in life sciences**, emphasizing that models should be both accurate and explainable (Rudin, 2019). By incorporating biochemical priors, such as energy conservation and positivity constraints, the neural-symbolic framework ensures biological realism while enabling scientific discovery rather than mere curve fitting.

### 4.5 Implications for metabolic engineering and disease modeling

The ability to derive **explicit energetic equations** from data has profound implications for both basic and applied science. In **metabolic engineering**, such models could inform strategies to optimize ATP production, minimize ROS generation, or fine-tune redox balance for biotechnological processes. By learning from measured metabolite trajectories, the framework could identify regulatory bottlenecks or emergent behaviors that classical flux balance analysis cannot capture.

In **disease contexts**, especially in neurodegeneration and cancer, where mitochondrial dysfunction is central, interpretable energetic models may elucidate how subtle shifts in energy distribution trigger pathological transitions. For example, altered NADH/ROS ratios have been linked to replicative stress in tumor cells and to DNA repair failure in neurodegenerative disorders (Chandel, 2015; Wang & Bogenhagen, 2006). A neural-symbolic system capable of identifying these thresholds could thus serve as a computational biomarker for **energy state-dependent disease progression**.

Moreover, the discovery of oscillatory energetic behavior may relate to **ultradian rhythms** and **metabolic clocks**, which coordinate transcriptional bursts and repair cycles (Feeney et al., 2016). These rhythmic features could represent an underlying “bioenergetic timing code” that governs when and how genetic information is accessed, a concept that could redefine our understanding of temporal control in molecular biology.

### 4.6 Toward quantum-informed generative bioenergetics

The framework presented here marks an early step toward what might be called **quantum-informed generative bioenergetics**, an emerging discipline that unites energy physics, AI, and molecular systems biology. In this vision, biological equations are not merely fitted to data but are *generated* from energetic principles that may include quantum coherence, entanglement, and tunneling phenomena within redox and proton transport chains (Lambert et al., 2013; Marais et al., 2018).

Generative AI models trained on high-resolution experimental data could extend this work to uncover new energy laws that span scales from molecular vibration to gene regulation. Such approaches may reveal how **quantum mechanical phenomena influence biological computation**, leading to predictive models that describe how the flow of energy gives rise to the flow of information, a concept central to understanding life as an energy-driven network.

### 4.7 Limitations and future directions

While the hybrid model achieved high accuracy and interpretability, several limitations remain. The simulated data were idealized and lacked the full biochemical complexity of living cells. Future work should incorporate **experimental time-series data** from mitochondrial or single-cell metabolomic measurements to validate and refine the derived equations.

Moreover, extending the framework to include **spatial diffusion, multi-compartment coupling**, and **quantum-inspired stochastic terms** could enable the modeling of real subcellular energy landscapes. Finally, integration with **multi-omics datasets**, including transcriptomics, proteomics, and metabolomics, would allow direct inference of gene-metabolism feedback loops, pushing the boundary toward fully **explainable AI-based systems biology**.

### 4.8 Summary and outlook

In conclusion, this work demonstrates that **interpretable machine learning**, when grounded in physical bioenergetics, can reveal the underlying logic of cellular decision-making. By uniting neural differential equations with symbolic discovery, we recover the mathematical syntax that links energy flow to genetic regulation.

The broader implication is that life may be best understood as a **computational expression of energy**, where ATP, NADH, and ROS act as both fuel and information carriers. As AI models grow increasingly quantum-informed and biologically aware, they may soon allow us to write, manipulate, and even design the energetic equations that sustain living systems.

## Supplementary Information

**SI Table 1.** Learned kinetic parameters.

| Parameter | Description | Learned Value |
| --- | --- | --- |
| k_trans | ATP–gene coupling | 0.722 |
| k_rep | Replication rate constant | 0.638 |
| k_repair | Repair energy coefficient | 0.726 |
| k_ros_decay | ROS decay constant | 0.114 |
| k_nadh_decay | NADH decay constant | 0.054 |

**SI Table 2.** Symbolic regression equations (top-ranked)

| Equation | R<sup>2</sup> | Description |
| --- | --- | --- |
| $G' = \alpha^1 ATP \cdot e^{-\beta_1 ATP} + \gamma^1 \sin(\delta^1 ATP)$ | 0.93 | Gene expression rate |
| $R' = \alpha^2 ATP^2 \cdot e^{-\beta_2 ATP}$ | 0.91 | DNA repair efficiency |
| $P' = \alpha^3 \left(\frac{NADH}{ROS}\right)^{\gamma^3}$ | 0.89 | Replication scaling with redox |

